# Climate change and the potential distribution of *Xylella fastidiosa* in Europe

**DOI:** 10.1101/289876

**Authors:** Martin Godefroid, Astrid Cruaud, Jean-Claude Streito, Jean-Yves Rasplus, Jean-Pierre Rossi

## Abstract

The bacterium *Xylella fastidiosa (Xf)* is a plant endophyte native to the Americas that causes worldwide concern. *Xf* has been recently detected in several regions outside its natural range including Europe. In that context, accurate estimates of its response to climate change are required to design cost-efficient and environment-friendly control strategies. In the present study, we collected data documenting the native and invaded ranges of the three main subspecies of *Xf: fastidiosa, pauca* and *multiplex*, as well as two strains of *Xf* subsp. *multiplex* recently detected in southern France (ST6 and ST7). We fitted bioclimatic species distribution models (SDMs) to forecast their potential geographic range and impact in Europe under current and future climate conditions. According to model predictions, the geographical range of *Xf* as presently reported in Europe is small compared to the large extent of suitable areas. The European regions most threatened by *Xf* encompass the Mediterranean coastal areas of Spain, Greece, Italy and France, the Atlantic coastal areas of France, Portugal and Spain as well as the south-western regions of Spain and lowlands in southern Italy. Potential distribution of the different subspecies / strains are contrasted but all are predicted to increase by 2050, which could threaten several of the most economically important wine-, olive- and fruit-producing regions of Europe, warranting the design of control strategies. Bioclimatic models also predict that the subspecies *multiplex* might represent a threat to most of Europe under current and future climate conditions. These results may serve as a basis for future design of a spatially informed European-scale integrated management strategy, including early detection surveys in plants and insect vectors, quarantine measures as well as agricultural practices.

## Introduction

The bacterium *Xylella fastidiosa (Xf)* is a plant endophyte native to the Americas, that develops in up to 300 plant species including ornamental and agricultural plants ^1^. In its native range, *Xf* is transmitted between plants by xylem-feeding insects belonging to several families of Hemiptera (Aphrophoridae, Cercopidae, Cicadellidae, Cicadidae and Clastopteridae) ^2^. *Xf* causes severe plant pathologies leading to huge economic losses ^3^, including the Pierce’s disease of grapevines PD: ^4^, the olive quick decline ^5^, the oak bacterial leaf scorch ^6^, the phony peach disease ^7^, the *Citrus* variegated chlorosis CVC: ^8^ and the almond leaf scorch ^9^. As *Xf* can colonize a large number of economically important plants including vine ^1^, its biology and the mechanisms of vector transmission have been extensively studied to design management strategies ^10^. On the basis of genetic data obtained with Multilocus Sequence Typing MLST: ^11,12^, *Xf* was subdivided into six subspecies (*fastidiosa, pauca, multiplex, sandyi, tashke* and *morus*). The subspecies were further characterized by different geographic origins, distributions and host preferences in the Americas ^13-15^. However, the status of the different subspecies is still a matter of debate ^16^ and only two, *fastidiosa* and *multiplex*, are formally considered valid names ^1,17^. *Xf* subsp. *fastidiosa* ^18^ occurs in North and Central America, where it causes, among others, the harmful PD and the almond leaf scorch (ALS). Genetic analyses suggest that this subspecies originated from southern parts of Central America ^19^. The subspecies *multiplex* is widely distributed in North America (from California to western Canada and from Florida to eastern Canada), where it was detected on a wide range of host plants (e.g. oak, elm, maple, almond, sycamore, *Prunus* sp., etc.) as well as in South America ^20,21^. The subspecies *pauca*, which causes severe diseases in *Citrus* (CVC) and coffee (Coffee Leaf Scorch: ^22^) in South and Central America, is thought to be native to South America ^23^. The subspecies *morus* recently proposed by Nunney et al ^24^, occurs in California and eastern USA, where it is associated to mulberry leaf scorch. *Xf* subsp. *sandyi*, responsible for oleander leaf scorch, is distributed in California ^12^, while the subspecies *tashke* was proposed by Randall et al ^25^ for a strain occurring on *Chitalpa tashkentensis* in New Mexico and Arizona. Overall, intraspecific entities of *Xf* display differences in host range suggesting that the radiation of *Xf* into multiple subspecies and strains is primarily associated to host specialization ^26^.

*Xylella fastidiosa* is now of worldwide concern as human-mediated dispersal of contaminated plant material has allowed the bacterium to spread outside its native range. In 2013, the CoDIRO strain (subsp. *pauca*) was detected on olive trees in southern part of the Apulia territory (Italy). Genetic analyses suggested that it was accidentally introduced from Costa Rica or Honduras with infected ornamental coffee plants ^5^. Since then, *Xf* subsp. *pauca* has spread northward and killed millions of olive trees in the Apulia territory, causing unprecedented socio-economic issues. During the period 2015-2017 several subspecies and strains were detected on *ca* 30 different host plants in Southern France (PACA region) and Corsica ^27^. According to this survey, the vast majority of plant samples were contaminated by two strains of *Xf* subsp. *multiplex* (referred here to as the French ST6 and ST7 strains). These strains are closely related to the Californian strains Dixon (ST6) and Griffin (ST7), belonging to the “almond group” of Nunney et al (2013) and were detected on numerous plant species though without evident specialization. To a lesser extent, other strains were detected in Southern France. Thus, the strain ST53 (Xf subsp. *pauca)* was detected on *Polygala myrtifolia* in Côte d’Azur (Menton) and on *Quercus ilex* in Corsica ^27^. Finally, recombinants strains (ST76, ST79 or not yet fully characterized) were detected in a few plant samples. In 2016, *Xf* subsp. *fastidiosa* was detected on rosemary and oleander plants overwintering in a nursery in Germany ^28^. In 2017, Spanish plant biosecurity agencies officially confirmed the detection of *Xf* strains belonging to the subspecies *multiplex, pauca* and *fastidiosa* on almond trees, grapevine, cherry and plums in western parts of the Iberian Peninsula and Balearic islands ^29^. Outside Europe, the detection of *Xf* was officially confirmed in Iran on almond trees and grapevines ^30^, in Turkey ^31^ as well as in Taiwan on grapevines ^32^. The severity of *Xf*-induced diseases has recently dramatically increased in several areas possibly due to global warming ^33^. Indeed, it has been demonstrated that cold winter temperatures might affect the survival of *Xf* in xylem vessels and allow plants to partly recover from *Xf*-induced diseases ‘cold curing phenomenon’: ^34,35^. For instance, Purcell ^34^ showed that grapevines with symptoms of PD recovered after multiple exposures to temperatures below −8°C during several hours. Further, Anas et al ^36^ suggested that areas experiencing more than 2 to 3 days with minimal temperature below −12.2 °C (or alternatively 4 to 5 days below −9.9 °C) should be considered at low risk for PD incidence, although these thresholds were considered too conservative by Lieth et al ^37^. There is no doubt that estimating the potential distribution of *Xf* under current and future climate conditions will contribute to design environment-friendly and cost-efficient management strategies ^38^. Several studies aimed to forecast the potential distribution of *Xf* in Europe ^39^ and/or all over the world ^40^. For instance, Hoddle et al ^40^ used the CLIMEX algorithm to forecast the worldwide potential severity of PD without accounting for potential future range shifts induced by global change. Bosso et al ^39^ fitted a Maxent model to forecast the potential distribution of *Xf* subsp. *pauca* under current and future climate conditions, and concluded that climate change would not affect its future distribution of *Xf*. However, Bosso et al ^39^ calibrated their model using solely the presence records from the invaded range in Italy, a practice that is known to increase omission errors ^41^.

Here, to assist in designing efficient survey as well as appropriate management strategies of *Xf* in Europe, we modeled the potential distribution of three of its subspecies: *fastidiosa, pauca* and *multiplex* under current and future climate conditions. For finest estimates we also modeled the potential distribution of the two strains of the subsp. *multiplex* that seem largely distributed in Southern France (i.e. the French ST6 and ST7). Finally, to go one step further, we estimated the severity of the Pierce’s Disease (caused by *Xf* subsp. *fastidiosa*) in European ecosystems based on US Pierce Disease intensity/occurrence maps provided by A. Purcell (available in Kamas et al ^42^ and in Anas et al ^36^).

## Material & Methods

### Distribution data

We collected occurrence data for subspecies *pauca* and *multiplex* in both their native and invaded ranges from the scientific literature, field surveys and public databases (Fig. 1). We also used data on the distribution of the strains ST6 and ST7 in France that were collected in 2015-2017 and stored in the French national database managed by the French Agency for Food, Environmental and Occupational Health & Safety (ANSES) (Fig. 1B). For the subspecies *fastidiosa*, we randomly generated 400 occurrences within its traditional range and assigned to each record a PD severity index (index 1: low severity; index 2: moderate severity; index 3: high severity) according to PD intensity maps provided by A. Purcell (available in Anas et al ^36^ and Kamas et al ^42^) (Fig. 1A). High severity regions comprise Florida, south of Alabama, Texas, Louisiana and Mississippi states as well as Los Angeles basin in California and coastal areas of South and North Carolina (Fig. 1A). Moderate severity areas comprise north of Alabama, Georgia, Texas, Louisiana and Mississippi states as well as some wine-producing regions of California (Napa valley, Sonoma valley, Santa Clara valley) where severe PD outbreaks occurred during the 20^th^ century even though it was an unusual phenomenon. Low severity regions encompass most of Virginia, Oklahoma, North Texas, Kentucky etc. as well as localities in Washington State ^34^. Pierce’s disease symptoms associated to the presence of *Xf* were recently detected in West Virginia ^33^, Oklahoma ^43^ and high elevation regions of Texas ^42^. The emergence of PD in these regions may, however, have been induced by the recent increase of temperatures occurring in the late 1990’s and in 2000’s. We deliberately considered these regions as low-risk areas because climate descriptors used in the present study represent average climate conditions for the 1970-2000 period (see below) and do not account for temperature changes that occurred at the very end of the 20^th^ century and the beginning of the 21^th^ century.

**Figure 1.**
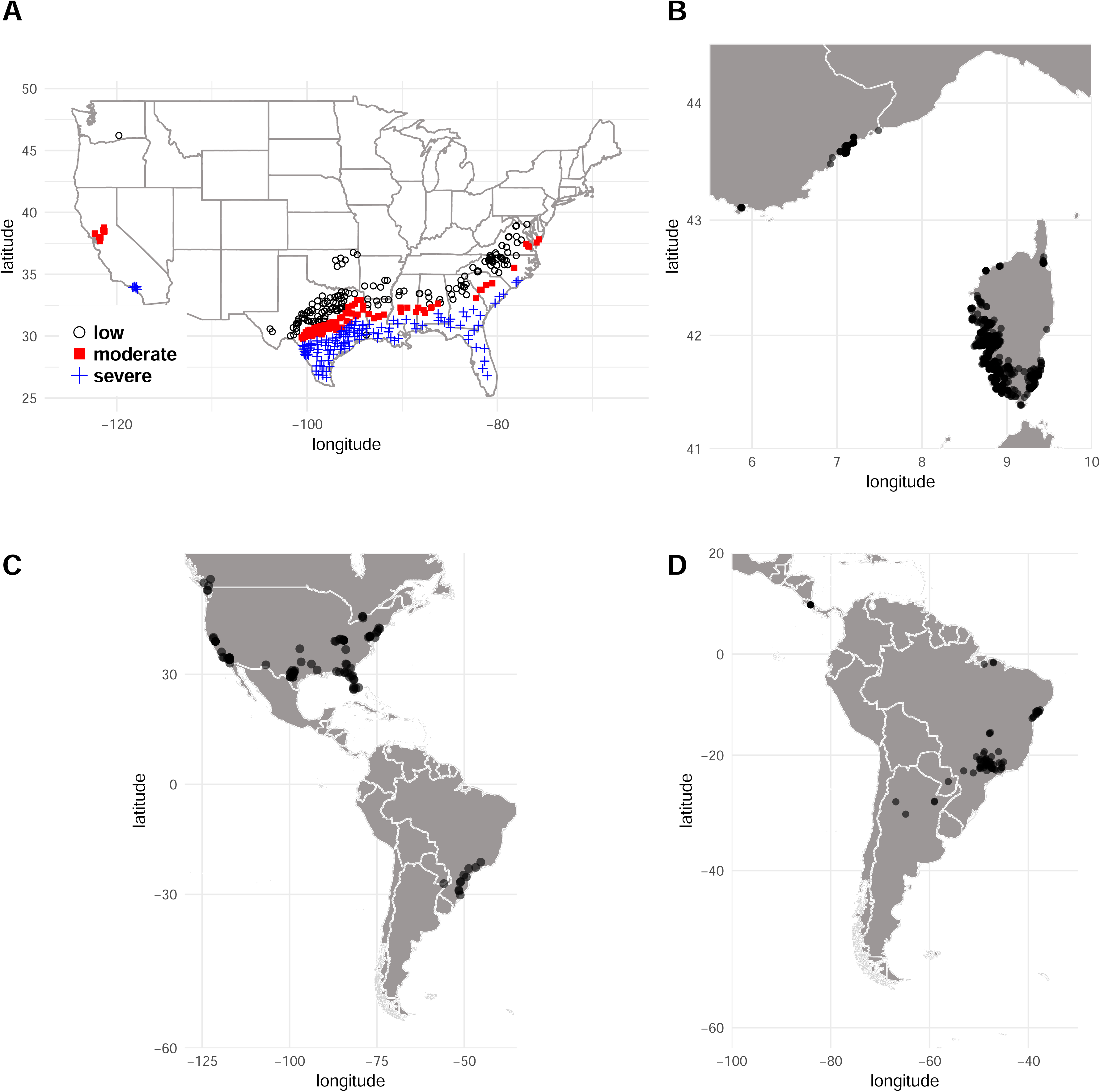
(A) Pierce’s disease (PD) severity map in the United States. Each locality is associated to a PD severity index (low, moderate or high severity) on the basis of the map available in Anas et al ^36^ and Kamas et al ^42^). (B) Occurrence records for the ST6 and ST7 strains in France. Occurrence records for (C) *Xylella fastidiosa* subsp. *pauca* and (D) *Xylella fastidiosa* subsp. *multiplex* in the Americas.

### Bioclimatic descriptors

We used a set of bioclimatic descriptors hosted in the Worldclim database bio1 to bio19 see ^44^. We used raster layers of 2.5-minute spatial resolution, which corresponds to about 4.5 km at the equator. The data come in the form of a raster map and represent the average climate conditions for the period 1970-2000. We estimated the future potential distributions of the different subspecies of *Xf* in 2050 and 2070 using the Model for Interdisciplinary Research on Climate version 5 MIROC5 ^45^. These predictions of future temperature and precipitation rank among the most reliable according to model evaluation procedures used in AR5 of IPCC ^46^. We used two different climatic datasets relative to the representative concentration pathways RCP4.5 and RCP8.5, which assume moderate and extreme global warming, respectively ^47^.

### Models

The PD severity index being an ordinal categorical variable (1<2<3) it was modeled using a cumulative link model (CLM) also called ordinal regression models, or proportional odds models ^48^. The CLM analyses the relationship between a set of independent variables (the climate descriptors) and an ordinal dependent variable consisting in the PD severity index. The CLM was fitted using a dataset corresponding to Pierce’s disease intensity in the US (Fig. 1A). It was then used to compute the spatial distribution of the probabilities that the different classes of index occur in Europe. CLM fit and predictions were carried out using the R software version 3.3.3 ^49^ and the R packages `MASS` ^50^ and `ordinal` ^51^.

The potential distribution of *Xf* subsp. *pauca, multiplex* and the French ST6/ST7 were estimated using species distribution modeling. Species distribution models establish mathematical species-environment relationships using presence records and environmental descriptors in order to assess the potential distribution of species or map the habitat suitability ^52^. We used two presence-only approaches namely Bioclim ^53,54^ and Domain ^55^. These algorithms are climatic envelope approaches. As such, they are based on presence records and do not make any assumptions about the absence of the organism under study. We selected these approaches because they are well suited to poorly documented species for which reliable absences are unavailable ^54,56^. In addition, we deliberately did not fit SDMs which rely on complex mathematical relationships among descriptors such as e.g. Maxent: ^57^ as we used only a few climate descriptors (see below).

Both Bioclim and Domain yield an index of habitat suitability that can be categorized to form a binary map (species presence vs. absence or suitable vs. unsuitable habitat). We used the lowest presence threshold (LPT) ^58^ i.e. the lower value of predicted climatic suitability associated to presence records. SDM fit and predictions were carried out using the R package `dismo` ^59^.

### Procedure to select climate predictors

The models were intentionally fitted using a limited number of ecologically relevant climate descriptors to avoid model over-parameterization, which is a recommended practice in the context of invasion risk assessment ^60^.

#### CLM

Although *Xf* geographical distribution appears to be primarily driven by minimum temperatures, the dynamics of the plant-pathogen-vector system is complex ^61^ and rainfall may impact the severity of the disease by affecting the bacterium pathogenicity or the intensity of the vection by insects.

The CLM was fitted using a set of climate variables that represent possible significant ecological stressors (maximum temperature of warmest month (bio5), minimum temperature of coldest month (bio6), mean temperature of warmest quarter (bio10), mean temperature of coldest quarter (bio11), precipitation of wettest quarter (bio16), precipitation of driest quarter (bio17), precipitation of warmest quarter (bio18) and precipitation of coldest quarter (bio19)). We used a stepwise model selection by AIC to identify the best performing variable subset, which finally comprised (bio10: mean temperature of warmest quarter, bio11: mean temperature of coldest quarter and bio18: precipitation of warmest quarter). Because the impact of precipitations upon PD severity pattern at large spatial scales is not well documented we performed the computations using the full model (bio10, bio11 and bio18) and a model comprising only the temperature predictors (bio10 and bio11).

#### SDM

A first step consisted into fitting and evaluating both Bioclim and Domain models using 10 different climate datasets combining maximal and minimal temperatures descriptors (bio5: maximum temperature of the warmest month; bio6: minimum temperature of the coldest month; bio10: mean temperature of the warmest quarter; bio11: mean temperature of the coldest quarter) (Table 1). We did not include descriptors of rainfall since we were interested in modeling distribution only (not severity) ^34^. At this stage, models were fitted using only the occurrences available in the native area of each subspecies or French occurrences of the ST6 and ST7 strains (Fig. 1B, C, D). This allowed us to evaluate the transferability of each model by calculating the proportion of actual presences in the invaded range correctly predicted using the LPT ^58^ i.e. model sensitivity. As a second evaluation of model transferability, we calculated the area under the curve of the receiver operator curve (AUC) from a dataset encompassing occurrences available in Europe as well as 10,000 pseudo-absences randomly generated across the European territory. For a given subspecies, the climatic dataset associated to models showing poor transferability were discarded from the study. The selected climate dataset were then used to fit Bioclim and Domain models using the occurrences available in both native and invaded ranges as recommended by various authors e.g. Broennimann and Guisan ^41^. The models were used to estimate the habitat suitability across Europe and binary maps were generated using the threshold detailed above. For each taxonomic unit we finally constructed a suitability map by averaging the predictions of models fitted with each selected set of climatic descriptors ensemble forecasting: ^62,63^.

**Table 1.** Measures of Bioclim and Domain models transferability calculated for different climate datasets. The area under the curve of the receiver operator curve (AUC) and the sensitivity of each model were calculated on the basis of occurrence records available in the European invaded range for *Xf* subsp. *pauca* and *multiplex*. The climate datasets leading to the best performing models were #1, #4 and #7 for subsp. *pauca* and #2, #6 and # 9 for subsp. *multiplex*. bio5: maximum temperature of the warmest month; bio6: minimum temperature of the coldest month; bio10: mean temperature of the warmest quarter; bio11: mean temperature of the coldest quarter.

| Dataset | Climate variables | Subspecies *pauca* |  |  |  | Subspecies *multiplex* |  |  |  |
| --- | --- | --- | --- | --- | --- | --- | --- | --- | --- |
|  |  | BIOCLIM |  | DOMAIN |  | BIOCLIM |  | DOMAIN |  |
|  |  | AUC | Sens. | AUC | Sens. | AUC | Sens. | AUC | Sens. |
| dataset #1 | bio6 | 0.88 | 1 | 0.9 | 1 | 0.686 | 0.999 | 0.759 | 0.999 |
| dataset #2 | bio11 | 0.44 | 0.04 | 0.78 | 0.04 | 0.918 | 0.999 | 0.924 | 0.999 |
| dataset #3 | bio6, bio11 | 0.45 | 0.04 | 0.79 | 0.12 | 0.742 | 0.999 | 0.847 | 0.999 |
| dataset #4 | bio5, bio6 | 0.96 | 1 | 0.96 | 1 | 0.839 | 0.999 | 0.797 | 0.999 |
| dataset #5 | bio5, bio11 | 0.5 | 0.04 | 0.92 | 0.16 | 0.838 | 0.999 | 0.801 | 0.999 |
| dataset #6 | bio10, bio11 | 0.5 | 0.04 | 0.91 | 0.16 | 0.884 | 0.999 | 0.965 | 0.999 |
| dataset #7 | bio6, bio10 | 0.95 | 1 | 0.95 | 1 | 0.867 | 0.999 | 0.904 | 0.999 |
| dataset #8 | bio5, bio6, bio11 | 0.5 | 0.04 | 0.92 | 0.16 | 0.843 | 0.999 | 0.803 | 0.999 |
| dataset #9 | bio6, bio10, bio11 | 0.5 | 0.04 | 0.91 | 0.16 | 0.871 | 0.999 | 0.933 | 0.999 |
| dataset #10 | bio5, bio6, bio10, bio11 | 0.5 | 0.04 | 0.92 | 0.16 | 0.853 | 0.999 | 0.819 | 0.999 |

### Refining species distribution models’ predictions

Some points in Europe may be associated to climate conditions that are not encountered within the range of conditions characterizing the set of reference points i.e. within the native and invaded areas. In such situations, using the species distribution models to predict habitat suitability in such novel habitats can be misleading ^64^. Elith et al ^64^ introduced the multivariate environmental similarity surface (MESS) to quantify how similar a point is to a reference set of points with respect to a set of predictor variables. Negative values of the MESS index indicate sites where at least one variable has a value lying outside the range of environments over the reference set. We computed the MESS index over Europe with reference to the occurrence dataset used to fit each species distribution model. We further restricted the model predictions to areas where the index was positive. We used a MESS index computed with the variable bio11 (mean temperature of coldest quarter) to refine the CLM predictions because the impact of winter temperatures on PD dynamics is very well documented ^34,35,65^. MESS computations were carried out using the R package `dismo` ^59^. All graphical outputs were produced using the R packages `ggplot2` ^66^ and `cowplot` ^67^.

## Results

### Pierce’s disease severity index

The stepwise-selected model comprised three bioclimatic descriptors: bio10: mean temperature of warmest quarter, bio11: mean temperature of coldest quarter and bio18: precipitation of warmest quarter. The variable contribution was highly significant in all cases (p<10^-3^) and the coefficients were −19.4 (bio10), 77.6 (bio11) and 3.5 (bio18). The resulting model was used to compute the values of the PD severity index across North America and Europe using climatic dataset corresponding to the period 1970-2000. In North America the MESS index revealed that areas north of 35 decimal degrees latitude were associated to strongly negative index (Fig. S1). In Europe, low index values were observed in north-eastern areas as well as in the Alps and the Pyrenees (Fig. S2). These areas were discarded from further interpretation and have been subsequently depicted in grey in the maps shown in Fig. 2. The Figs 2A and S3 show the model predictions for the three levels of severity in Europe and North America for the period 1970-2000 respectively.

**Figure 2.**
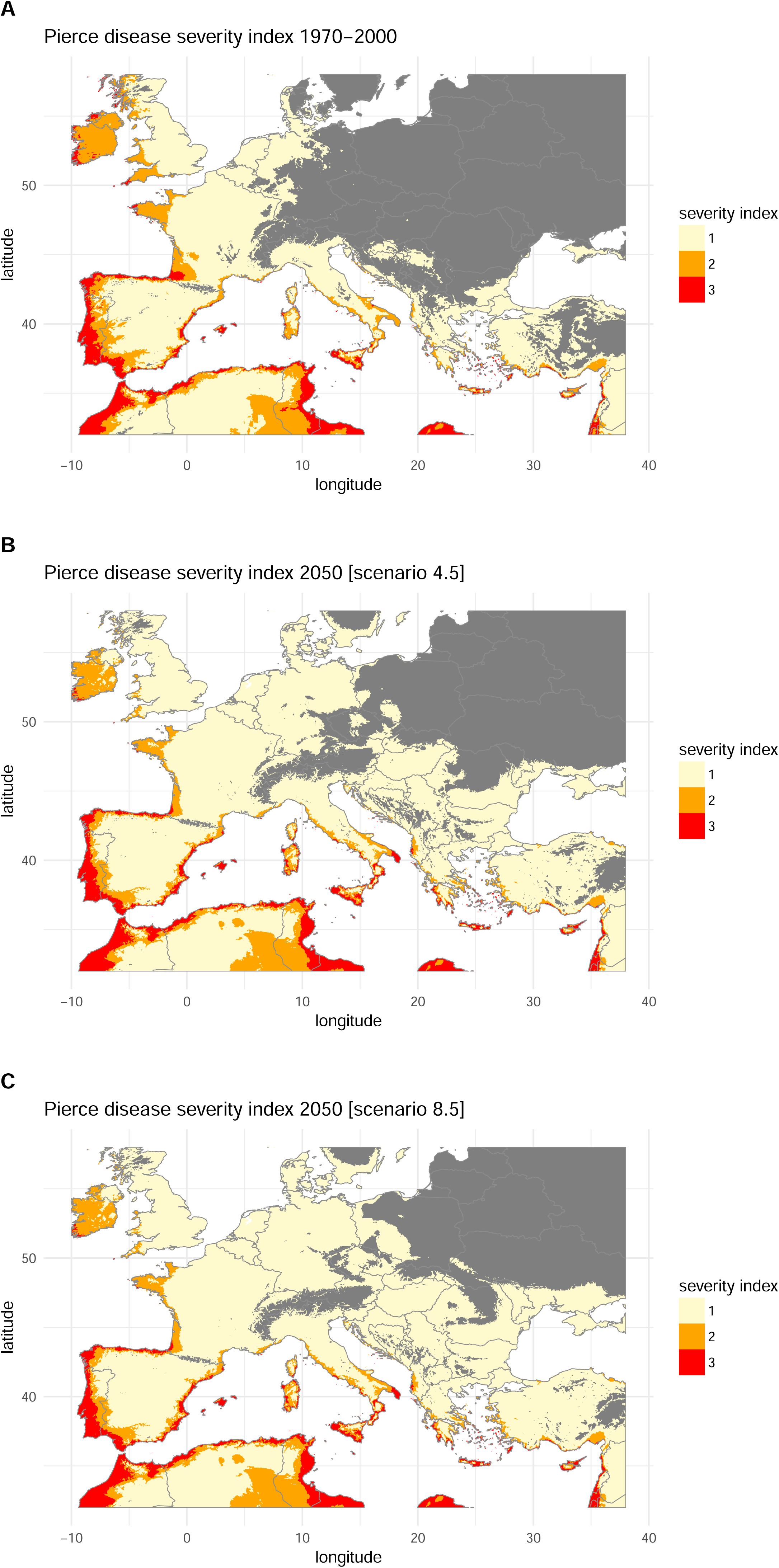
Predicted potential severity of Pierce’s disease in Europe under current and future climate conditions obtained from a cumulative link model (CLM). 1 = low severity, 2 = moderate severity, 3 = high severity. Current climate conditions are average temperature for the period 1970-2000 extracted from the Worldclim database ^44^. Future climate estimates were obtained from the MIROC5 global climate model (scenarios 4.5 and 8.5). A: Predicted PD severity index for the period 1970-2000. B: Predicted PD severity index in 2050 with the scenarios RCP4.5. C: Predicted PD severity index in 2050 with the scenarios RCP8.5. Areas associated to climate conditions that are not met within the range of conditions characterizing the set of reference points in the native range (i.e. MESS index < 0 see material and methods) are shown in grey.

The CLM predicted a risk of moderate to highly severe PD in multiple lowlands and coastal areas of the Mediterranean regions (Spain, Italy, Balearic islands and North Africa) as well as along the Atlantic coasts of France, Northern Spain and Portugal (Fig. 2A). A low to moderate severity of PD was also predicted in the Atlantic costs of France, lowlands of northern Italy, and central Spain. High severity was predicted in Sicilia (Italy), and long both the Atlantic and Mediterranean coasts of Spain.

Using the model to estimate the PD severity index according to different climate change scenarios led to the maps displayed in Fig. 2B and 2C for 2050 and Fig. S4 B and C for 2070. In each case, the MESS index was recomputed on the basis of the present and future climate conditions. Estimations for 2050 indicate an increased PD severity in south Italy, Corsica and Sardinia either with the concentration pathways RCP4.5 or the RCP8.5 (Fig. 2B, and 2C). The estimates for 2070 are pretty much similar (Fig. S4).

The CLM fitted with only bio10 and bio11 led to the results showed in Fig. S5 and S6. The main differences are that the Atlantic coasts of France, Ireland and west England are associated to severity index of 1 whereas the model including bio18 predicted a severity of 2 (Fig. S5). A similar pattern is observed for the predictions in 2070 (Figs S4 and S6).

### Potential distribution of Xf subspecies pauca and multiplex

Both Bioclim and Domain fitted using climate datasets #1, #4 and #7 yielded high transferability measures (AUC superior to 0.85 and sensitivity = 1) in the case of the subsp. *pauca* (Table 1). These datasets were therefore retained for further analyses. For the subsp. *multiplex* we selected 3 climate datasets (#2, #6 and #9) associated to AUC values >0.85 and a sensibility of 0.999 (Table 1).

Regarding the subspecies *pauca*, the models showed that climatically suitable environments in Europe only correspond to small well-delimited areas in South Portugal and Spain, Balearic Islands, Sicilia and North Africa (Fig. 3A). There was a marked agreement between the models that all indicated very favorable environment in these areas (Fig. 3D). It is worth noting that the areas associated to negative MESS index values are very large (shaded areas in maps; i.e. areas experiencing climate conditions absent from the dataset used to calibrate the model and for which no prediction was made). This conveys the fact the subsp. *pauca* originates from South America (Fig. 1D) and is associated to tropical environments. Suitable environments in Europe are restricted to warmest environments around the Mediterranean Sea. The models predict changes in the location of suitable areas in Europe by 2050 (Fig. 3B-F). Areas at risk would include northern coast of Spain, south France and Tyrrhenian coast of Italy. There is no marked differences according to the scenario examined (Fig. 3) and a similar pattern is expected in 2070 (Fig. S7) except for Italy where climate conditions may become unfavorable according to scenario 8.5 (Fig S7F)

**Figure 3.**
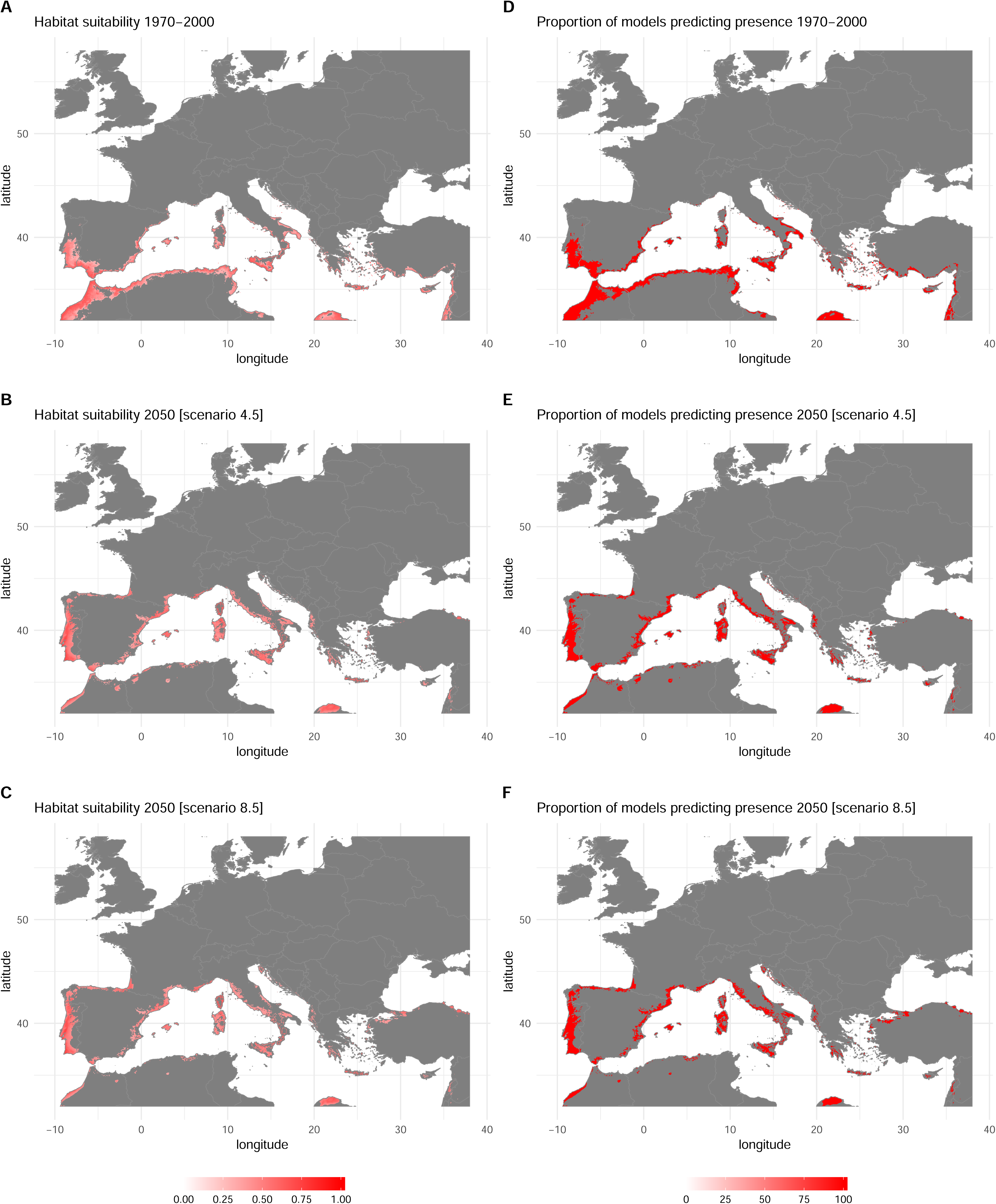
Predicted potential distribution of *Xf* subsp. *pauca* in Europe under current and future climate conditions obtained by fitting Bioclim and Domain models. Current climate conditions are average temperatures for the period 1970-2000 extracted from the Worldclim database. Future climate estimates were obtained from the MIROC5 global climate model (scenarios 4.5 and 8.5). A: Habitat suitability for the period 1970-2000. B: Habitat suitability in 2050 for the scenario RCP4.5. C: Habitat suitability in 2050 for the scenario RCP8.5. D: Proportion of models predicting presence for the period 1970-2000. E: Proportion of models predicting presence in 2050 for the scenario RCP4.5. F: Proportion of models predicting presence in 2050 for the scenario RCP8.5. Maps A, B, C were obtained by averaging (ensemble forecasting) of the outputs of the models Bioclim and Domain run with 3 different climate datasets (see details in Table 1). Maps D, E, F were obtained by averaging the presence/absence maps derived from habitat suitability using the lowest presence threshold. Areas associated to climate conditions that are not met within the range of conditions characterizing the set of reference points in the native range (i.e. MESS index < 0 see material and methods) are shown in grey.

The situation is different for the subspecies *multiplex* which is natively distributed across North America (Fig. 1C) and for which the models depicted suitable conditions in most of Europe except high-elevation areas and cold northern regions (Fig. 4). The expected impacts of climate change are limited and mostly concern South Spain where the conditions are expected to become unfavorable by 2050 and North part of Europe that are predicted to become more favorable by 2050 (Fig. 4 B to F) and 2070 (Fig. S8). The potential distribution of the French strains ST6 and ST7 is localized to Mediterranean areas (Corsica, Sardinia, Sicilia and coastal areas of Italy and France) (Fig. 5A, D). Suitable conditions are also present in the Atlantic coasts of Portugal and in South West France. A shift in distribution is expected to occur by 2050 (Fig. 5B to F and Fig. S9). Favorable conditions are expected to extend northward while areas currently suitable such as South Western France are expected to become unfavorable. All models indicate that the Spanish Atlantic coast (Galicia, Asturias, Cantabria and Basque country) is expected to become climatically suitable by 2050.

**Figure 4.**
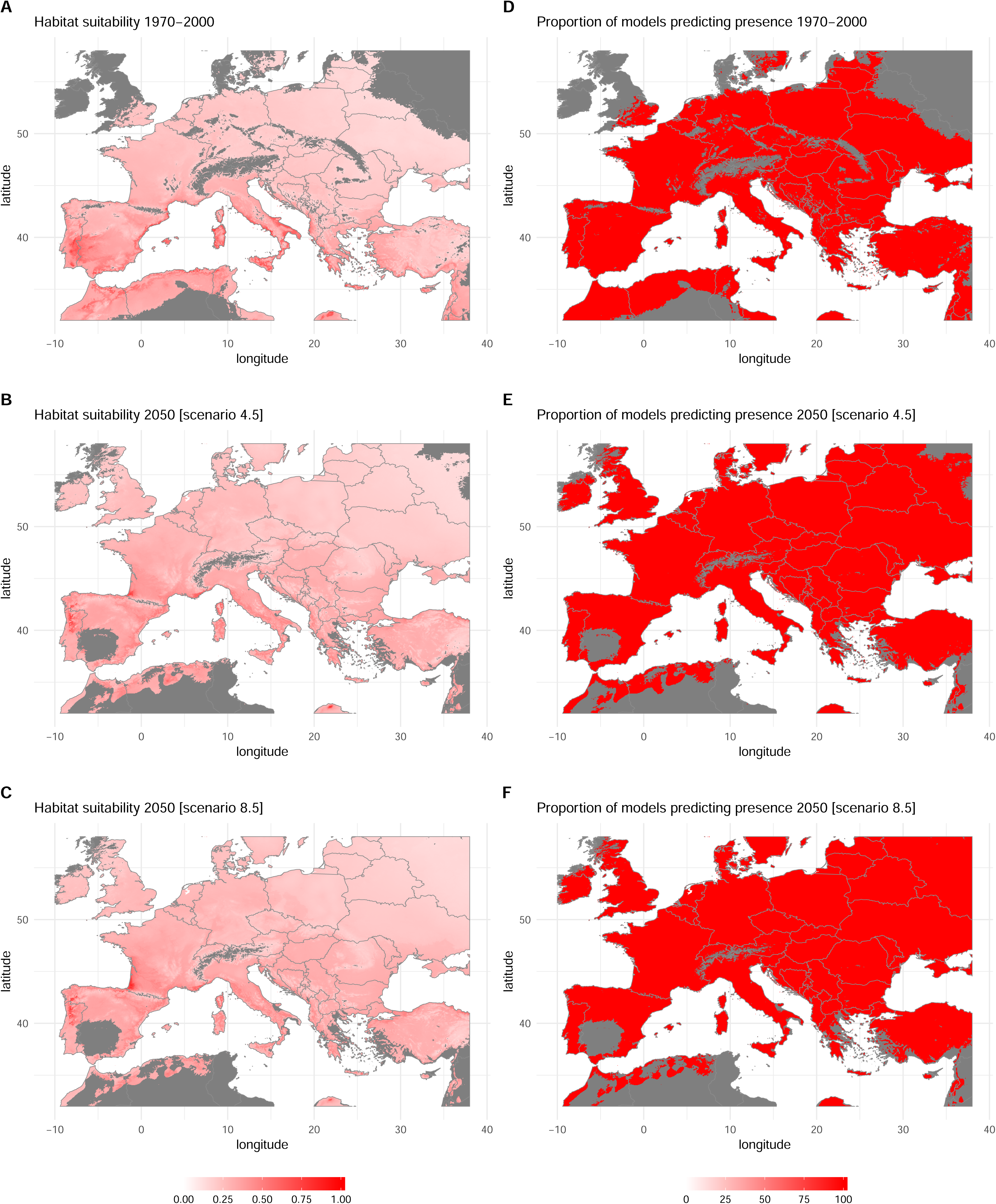
Predicted potential distribution of *Xf* subsp. *multiplex* in Europe under current and future climate conditions obtained by fitting Bioclim and Domain models. Current climate conditions are average temperatures for the period 1970-2000 extracted from the Worldclim database. Future climate estimates were obtained from the MIROC5 global climate model (scenarios 4.5 and 8.5). A: Habitat suitability for the period 1970-2000. B: Habitat suitability in 2050 for the scenario RCP4.5. C: Habitat suitability in 2050 for the scenario RCP8.5. D: Proportion of models predicting presence for the period 1970-2000. E: Proportion of models predicting presence in 2050 for the scenario RCP4.5. F: Proportion of models predicting presence in 2050 for the scenario RCP8.5. Maps A, B, C were obtained by averaging (ensemble forecasting) of the outputs of the models Bioclim and Domain run with 3 different climate datasets (see details in Table 1). Maps D, E, F were obtained by averaging the presence/absence maps derived from habitat suitability using the lowest presence threshold. Areas associated to climate conditions that are not met within the range of conditions characterizing the set of reference points in the native range (i.e. MESS index < 0 see material and methods) are shown in grey.

**Figure 5.**
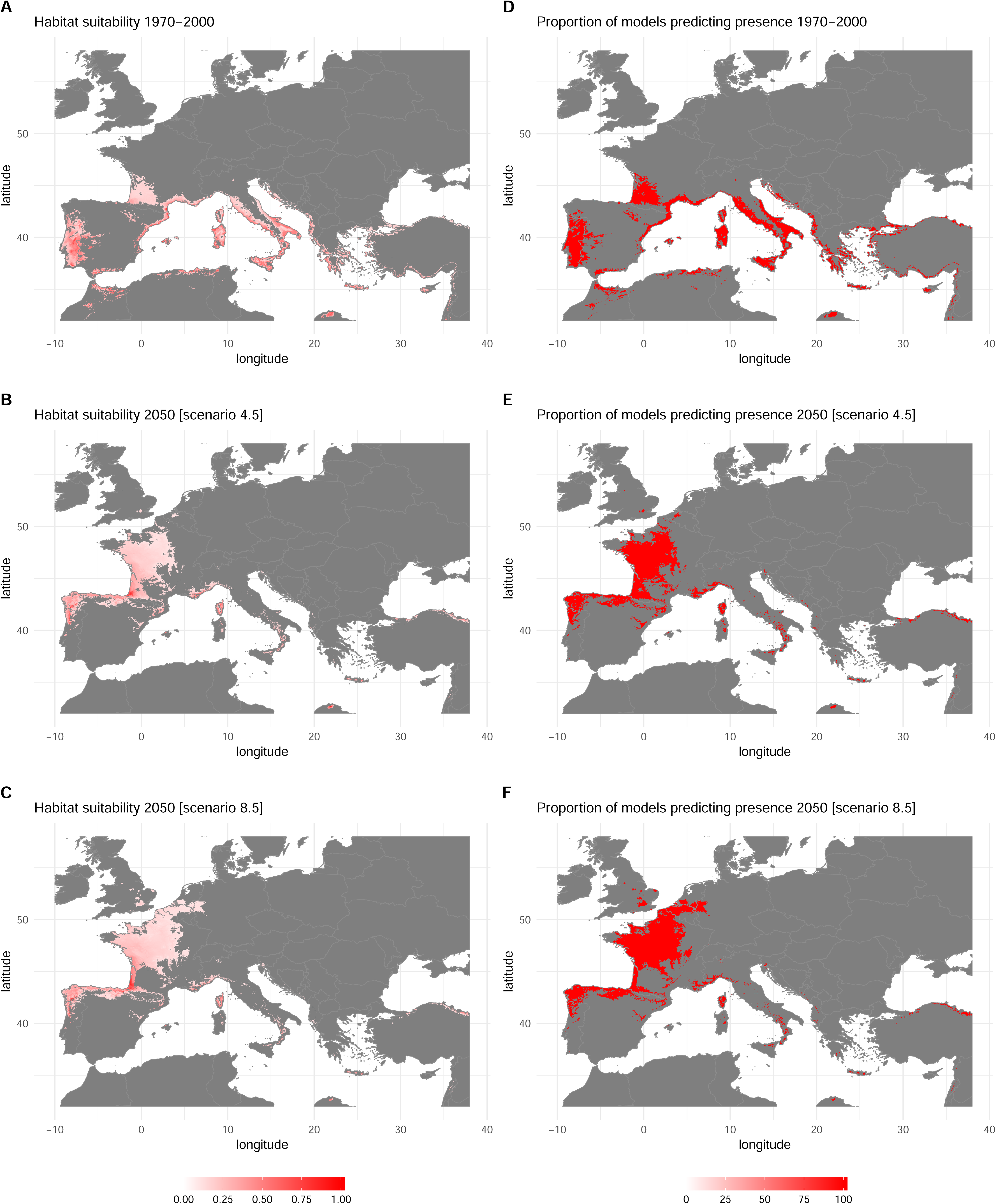
Predicted potential distribution of the French strains ST6 and ST7 (*Xf* subsp. *multiplex*) in Europe under current and future climate conditions obtained by fitting Bioclim and Domain models. Current climate conditions are average temperatures for the period 1970-2000 extracted from the Worldclim database. Future climate estimates were obtained from the MIROC5 global climate model (scenarios 4.5 and 8.5). A: Habitat suitability for the period 1970-2000. B: Habitat suitability in 2050 for the scenario RCP4.5. C: Habitat suitability in 2050 for the scenario RCP8.5. D: Proportion of models predicting presence for the period 1970-2000. E: Proportion of models predicting presence in 2050 for the scenario RCP4.5. F: Proportion of models predicting presence in 2050 for the scenario RCP8.5. Maps A, B, C were obtained by averaging (ensemble forecasting) of the outputs of the models Bioclim and Domain run with 3 different climate datasets (see details in Table 1). Maps D, E, F were obtained by averaging the presence/absence maps derived from habitat suitability using the lowest presence threshold. Areas associated to climate conditions that are not met within the range of conditions characterizing the set of reference points in the native range (i.e. MESS index < 0 see material and methods) are shown in grey.

## Discussion

### Geographical distribution and possible impacts in Europe

In a rapidly changing world, the design of pest control strategies (e.g. early detection surveys and planning of phytosanitary measures) should ideally rely on accurate estimates of the potential distribution and/or impact of pest species as well as their responses to climate change ^38^. In the present study, bioclimatic models predicted that a large part of the Mediterranean lowlands and Atlantic coastal areas are seriously threatened by *Xf* subsp. *fastidiosa, multiplex* and *pauca*. A low to moderate impact is also expected in northern and eastern regions of Europe (North-eastern France, Belgium, the Netherlands, Germany, Scandinavia, the Baltic region, Poland, Austria, Switzerland, etc.) that experience lower minimal temperature in winter but may nevertheless presumably host *Xf* subsp. *multiplex*.

Models display good evaluation measures and predict moderate to high climatic suitability in all European areas where symptomatic plants are currently infected by the subspecies *fastidiosa, pauca* or *multiplex* (e.g. Balearic Islands, lowlands of Corsica island, south-eastern France and the Apulia region). This suggests that risk maps provided in the present study are reliable for the design of surveys, including ‘spy insects’ survey ^68,69^. They may also be helpful to anticipate the spread of the different subspecies and provide guidance on which areas should be targeted for an analysis of local communities of potential vectors and host plants to design management strategies and research projects. Our results show that the subspecies/strains of *Xf* studied here might significantly expand in the near future, irrespective of climate change. For example, the ST6 and ST7 strains (subsp. *multiplex*) present in Corsica and southern France have a large potential for expansion, whose dynamics actually depends more on plant exchanges and disease management than on climate suitability *per se*. The subspecies *multiplex* is associated to economically important plants such as almonds and olives ^26^ but may also colonize multiple ornamental plants. Its present potential distribution in Europe extends far beyond areas where the subspecies has been reported and comprises Portugal, Italy, and both South and South-western France suggesting possible important economical losses.

The subspecies *fastidiosa*, which has been currently reported from a limited number of localities, could encounter favorable climate conditions in various areas of Europe. The model estimation of areas with a risk of PD highlights strategic wine-growing areas in different countries. Notably, the present estimates of the potential impact of the *subsp. fastidiosa* are consistent with the risk maps provided by Hoddle et al^40^ and A. Purcell (available in Anas et al ^36^). The case of subsp. *pauca* is somewhat different. Most of the European occurrences are known from southern Italy and the Balearic Islands and the potential distribution of this subspecies appears to be limited. This is not surprising given that *Xf* subsp. *pauca* is native from South America and occupies a climatic niche that mostly corresponds to areas located around the Mediterranean basin. Nevertheless, southern Spain, Portugal, Sicilia and North Africa that are areas where growing olive trees is multisecular offer suitable conditions, which potentially implies huge socio-economic impacts. One factor that proved to be critical for some diseases is the distribution/availability of vectors and hosts. Here, none of these factors is limiting since *Xf* is capable of colonizing a vast array of plants present in Europe and *Philaenus spumarius*, the only European vector identified so far ^70,71^, occurs across the whole continent ^69^.

Because we used the MESS index to discard regions experiencing climate conditions absent from the dataset used to calibrate the model, our estimations of potential distributions are conservative. The CLM showed a positive effect of higher temperatures during the coldest quarter (variable bio11 associated to a positive coefficient) on the severity index which may indicate to a lower “cold curing” effect ^34,65^. Absence of estimation of the potential distribution of *Xf* (all subspecies) or of the PD severity index (*Xf* subsp. *fastidiosa*) (i.e. shaded areas of the maps) does not mean that the bacterium is unable to develop but rather that evaluating the risk is difficult. For example, the potential impact of *Xf* in areas experiencing extremely high temperatures in summer (e.g. southern and central Spain) remains largely uncertain as the impact of extreme heat on *Xf* and on the behavior of insect vectors is still poorly known ^61^. We report a negative coefficient for the variable bio10 (mean temperature of the warmest quarter) suggesting that PD severity would be negatively related to high temperatures during summer. Although warm spring and summer temperatures enhance multiplication of *Xf* in plants, it has been showed that *Xf* populations decrease in grapevines exposed to temperatures above 37°C ^35^. As southern and central Spain frequently experience temperatures above 40°C in summer, further field and laboratory experiments are required to improve our estimation of the potential impact of *Xf* in these regions. Another point requiring clarification is the effect of precipitation during the warmest quarter that appears to be significant in the CLM. Precipitation may have direct impacts on the dynamics of the relationship between *Xf* and its host as well as indirect effects through the relationships with the insect vectors.

### Climate change and possible range shifts

Our results clearly indicate that climate change may strongly impact the distribution of *Xf* in Europe. Indeed, as “cold curing” appears to be the main mechanism explaining the lower impact of *Xf* in colder regions, an increase of winter temperatures should make these regions more suitable for *Xf* in the next decades ^34,35,65^. We report possible changes in the potential distribution of the subspecies *multiplex* with a northward expansion by 2070. The potential distribution of the French strains ST6 and ST7 is even more impacted with a gradual shift of suitable areas from Southern France, Italy and Portugal towards Northern France, Belgium, Netherlands and South England. The suitable areas for *Xf* subsp. *pauca* are expected to slightly extend to the Mediterranean coastal areas of Spain and France. The expected impact of climate change on the severity index of the PD is less marked and mostly correspond to an increased PD severity in south Italy, Corsica and Sardinia.

Overall, these results obtained on the different subspecies clearly indicate that climate change will alter areas at risk for invasion by *Xf* in Europe. Given that both the concentration pathways RCP4.5 and RCP8.5 led to concordant predictions, it appears sound to expect such changes even if the global warming is kept to a moderate level. SDMs showed that the subspecies *multiplex* displays a wider temperature tolerance and could threaten most of the European continent now and in the future. This is not surprising as this subspecies infects elms in regions of Canada characterized by low winter temperatures ^72^. This broad tolerance to cold is not known for other subspecies and it is still unclear whether realized niche divergence among subspecies reflects inherent differences in thermal tolerances or rather host-pathogen interactions as it was observed for *Ralstonia solanacearum* ^73^. More investigations would help a better understanding of the effect of temperatures on the different strains of *Xf*. It is noteworthy that both present and future distributions show several areas of potential co-occurrence. This may have important implications as it may increase the risk of intersubspecific homologous recombination (IHR).

### Limits and opportunities for risk assessment

Maps of habitat suitability and their declination with regards to future climate conditions should be guardedly interpreted as they are derived from correlative tools that depict the *realized* niche of species i.e. a subset of the *fundamental* environmental tolerances constrained by biotic interactions and dispersal limits ^74^. In addition, we cannot rule out the possibility that this study overestimates the potential distribution of the subspecies *pauca* and *multiplex*, and of the French strains ST6 and ST7 under future climate conditions. Indeed, time-periods associated to occurrences and climate descriptors dataset do not perfectly overlap. The models were fitted with climate descriptors that represent average climate conditions for the 1970-2000 period, while most presence records were collected after 2000 in a period characterized by milder winter temperatures. Moreover, as we deliberately fitted simple climate-envelope approaches such as Bioclim and Domain based on few climate descriptors to avoid model over-parameterization and/or extrapolation and enhance model transferability, we cannot exclude that bioclimatic models presented in the study do not capture the entire range of environmental tolerances and do not depict the complexity of the climatic niche of *Xf* as well as potential interactions between climate descriptors. Better models hence better risk assessment could be obtained if the amount of occurrence data could be increased. True absences i.e. locations where the environmental conditions are unsuitable for *Xf* to survive, would be particularly precious because they would allow using powerful approaches such as the generalized linear model ^52^. Finally, the possible adaptation of the subspecies of *Xf* to environmental constraints met in European ecosystems is another important and unknown factor of uncertainties. For example, the potential of recombinant strains to adapt should be addressed in the near future.

Finally, it is worth noting that bioclimatic models predict climatic suitability of a geographic region rather than a proper risk of *Xf*-induced disease incidence. To predict the proper severity of *Xf*-induced diseases in a given locality, statistical models should account for many additional factors playing a role in *Xf* epidemiology, including e.g. microclimate conditions, inter-annual climate variability, landscape structure and the spatio-temporal structure of the community of potential vectors. Although recent entomological studies identified the meadow spittlebug *P. spumarius* as the main vector of *Xf* in Italy ^70,71^, a better knowledge of all European vectors capable of transmitting *Xf* from plant to plant as well as their ecological characteristics (geographic range, efficiency in *Xf* transmission, demography, overwintering stage, etc.) is needed ^75^. In this context, our estimates could allow to design cost-efficient vector surveys, with priority given to geographic regions predicted as highly climatically suitable for *Xf*. The study by Cruaud et al ^69^ provides a good insight into how species distribution modeling and DNA sequencing approaches may be combined for an accurate monitoring of the range of *Xf* and its vectors in Europe. We believe that bioclimatic models are promising tools to help designing research experiments, control strategies as well as political decisions at the European scale.

## Conclusions/highlights

Species distribution models all indicate that the geographical range of *Xf* as presently reported in Europe is small compared to the large extent of suitable areas. This is true for all studied subspecies of *Xf* although the subspecies *pauca* appears to have a smaller potential range possibly because of its Neotropical origin. Although caution is needed in interpreting spatial projections because uncertainties in future climate conditions themselves and because uncertainties associated to the predictions of the species distribution models are difficult to assess, we showed that climate change will probably affect the future distribution of the bacterium by 2050 and then 2070. Last but not least, *Xf* has a certain potential to adapt for the specific climate and biotic interactions (hosts, vectors) present in Europe. This potential is unknown but could nevertheless lead to marked divergence between its future geographical distribution and the picture we have of it today. However, our current knowledge allows proposing different research avenues to better understand and anticipate the possible expansion of *Xf* in Europe. European areas at risk encompass diversified (sub)natural ecosystems as well as agro-ecosystems in which an important research effort should be made to decipher the host plants – insect vectors – bacterium interactions ^76^. Only in this way could we develop an appropriate and efficient strategy to deal with *Xf* in the coming years.

## Acknowledgements

We thank Pauline De Jerphanion (ANSES) manager of the French national database of *Xylella fastidiosa* in France as well as the DGAL and the SRAL or feeding that database. We also thank Christian Lannou (INRA, SPE, France) for his interest and support during the course of the project. This work was funded by grants from the SPE department of the INRA (National Agronomic Institute). The funders had no role in study design, data collection and analysis, decision to publish, or preparation of the manuscript. JPR thanks the SNCF (French national railway company) whose recurrent delays allowed him to deeply meditate on the results reported in here.

**Supplementary Information accompanies this paper and are available as a** pdf file (Godefroid_etal_Xylella_fastidiosa_Europe_suppl_mat.pdf)

