## Supplementary Materials for "Climate change and the potential distribution of *Xylella fastidiosa* in Europe"

CBGP, INRA, CIRAD, IRD, Montpellier SupAgro, Univ. Montpellier, Montpellier, France

[1]Present address: Fisheries & Oceans Canada, Pacific Biological Station, Nanaimo, B.C

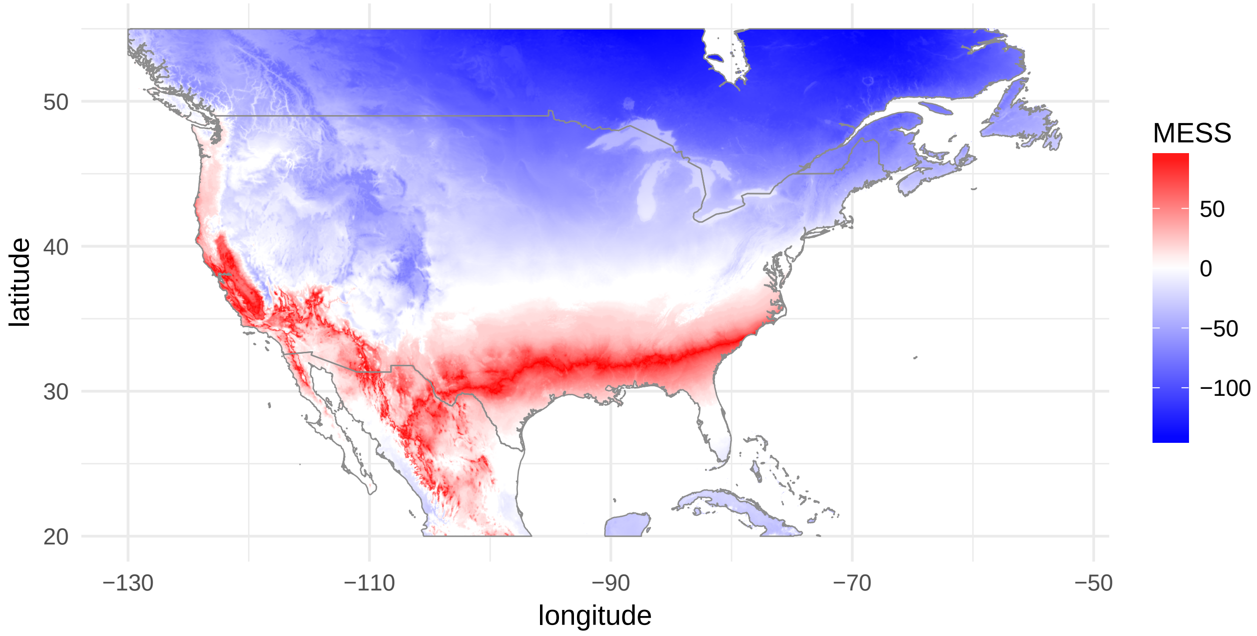

**Figure S1** – Multivariate environmental similarity surface (MESS)[1] index computed for North America with reference to a set of points corresponding to localities where *Xylella fastidiosa fastidiosa* was reported in the USA (see Figure 1A). The predictor variable is the mean temperature of coldest quarter (bio11) considered to represent a very strong constraint on the distribution of the bacterium. Legend: blue, negative; white, around zero; red, negative; with more intense colors indicating more extreme values. The climate data were downloaded from the worldclim version2 web site `http://worldclim.org/version2`.[2]

[1] Elith J., Kearney M. and Phillips S. 2010. The art of modelling range-shifting species. Methods Ecol Evol 1:330-342.

[2] Fick S.E. and Hijmans R.J. 2017. WorldClim 2: new 1km spatial resolution climate surfaces for global land areas. Int J Climatol 37:4302-4315.

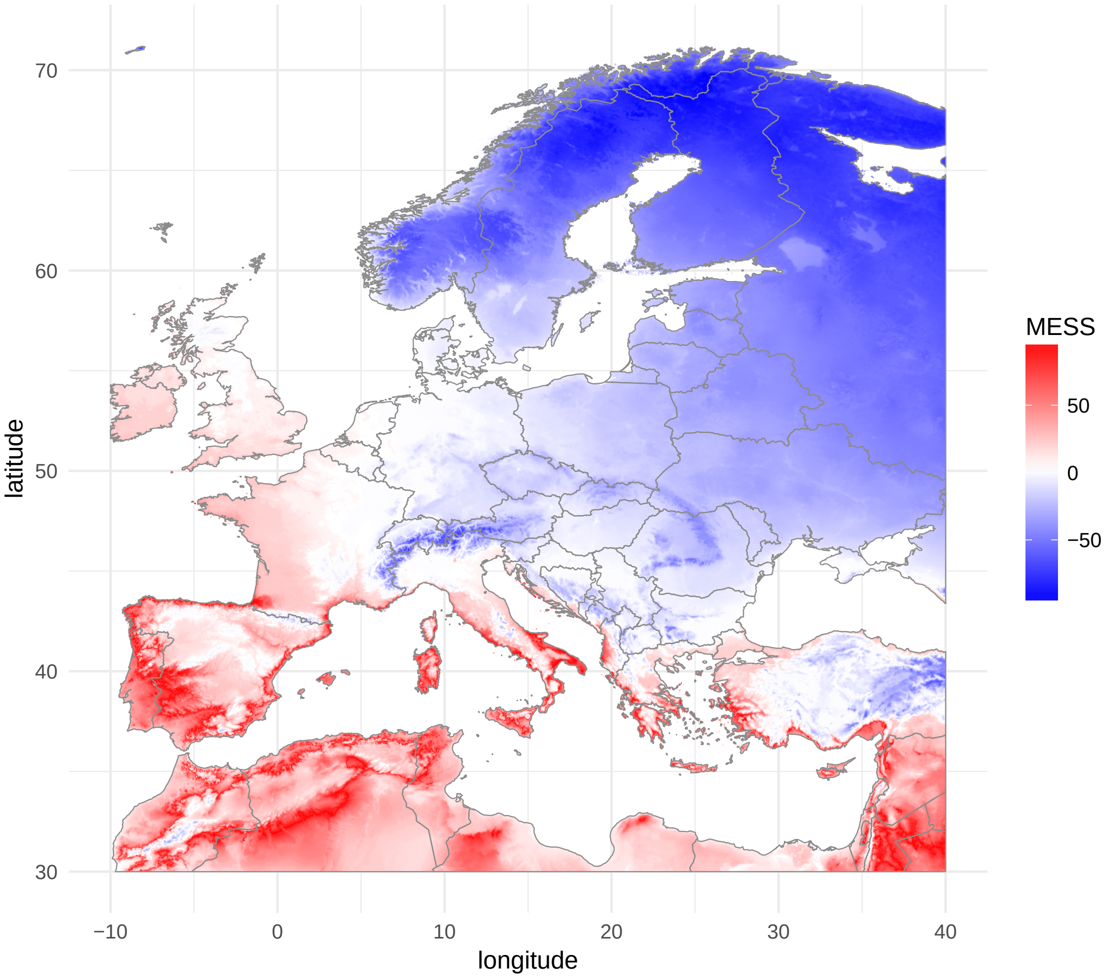

**Figure S2** – Multivariate environmental similarity surface (MESS) index computed for Europe with reference to a set of points corresponding to localities where *Xylella fastidiosa fastidiosa* was reported in the USA (see Figure 1A). The predictor variable is the mean temperature of coldest quarter (bio11) considered to represent a very strong constraint on the distribution of the bacterium. Legend: blue, negative; white, around zero; red, negative; with more intense colors indicating more extreme values. The climate data were downloaded from the worldclim version2 web site `http://worldclim.org/version2`.

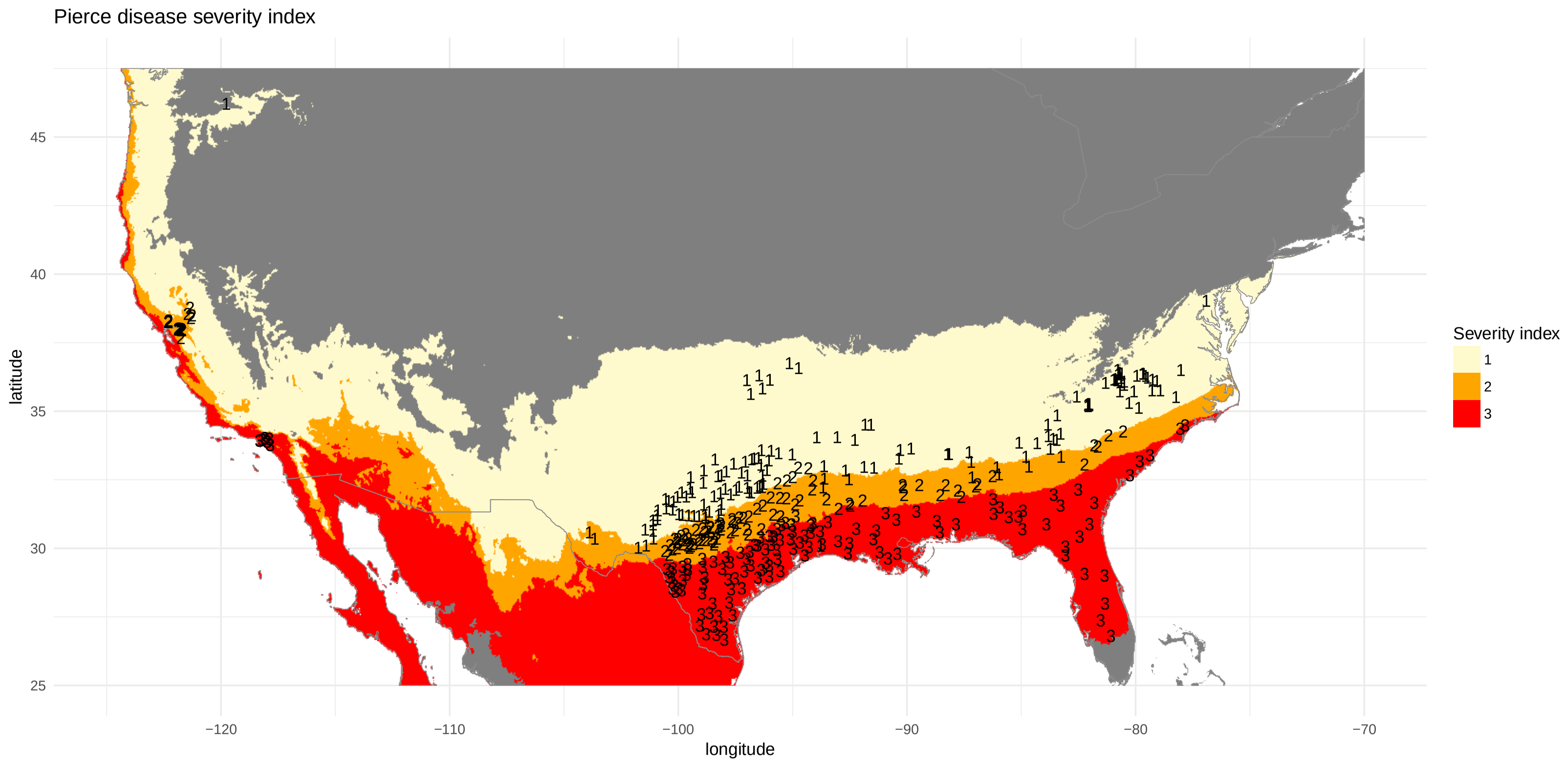

**Figure S3** – Estimation of the Pierce severity index in North America. The index is an ordinal variable with 3 modalities 1: low severity; 2: moderate severity and 3: high severity defined by Purcell[1] (see Figure 1A). Numbers correspond to reference values. A cumulative link model was fitted to link these values to 3 bioclimatic predictors, the mean temperature of warmest quarter, the mean temperature of coldest quarter and the precipitation of warmest quarter referred to as bio10, bio11 and bio18 in the worldclim database `http://worldclim.org/version2`. The model was used to compute the index across North America. The predictions associated to localities with a negative MESS index were discarded and depicted in grey.

[1] Anas O., Harrison U.J., Brannen P.M. and Sutton, T.B. 2008. Effect of warming winter temperatures on the severity of Pierce's disease in the Appalachian Mountains and Piedmont of the Southeastern United States. Plant Health Prog 1:450-459.

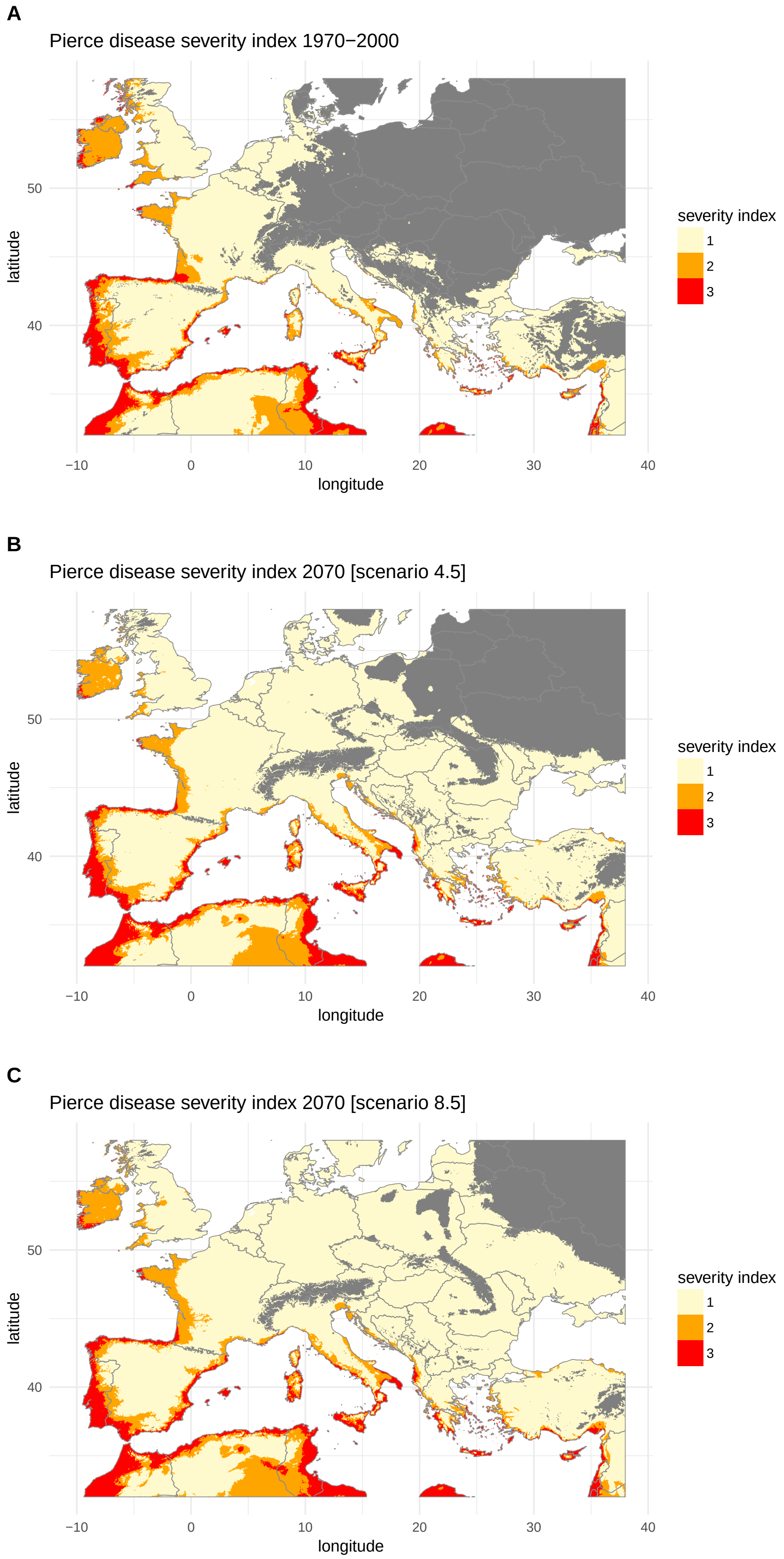

**Figure S4** – Predicted potential severity of Pierce's disease in Europe under current and future climate conditions obtained from a cumulative link model (CLM). 1 = low severity, 2 = moderate severity, 3 = high severity. Current climate conditions are average temperature for the period 1970-2000 extracted from the Worldclim database (`http://worldclim.org/version2`). Future climate estimates were obtained from the MIROC5 global climate model (scenarios 4.5 and 8.5). A: Predicted PD severity index for the period 1970-2000. B: Predicted PD severity index in 2070 with the scenarios RCP4.5. C: Predicted PD severity index in 2070 with the scenarios RCP8.5. The predictions associated to localities with a negative MESS index were discarded and depicted in grey.

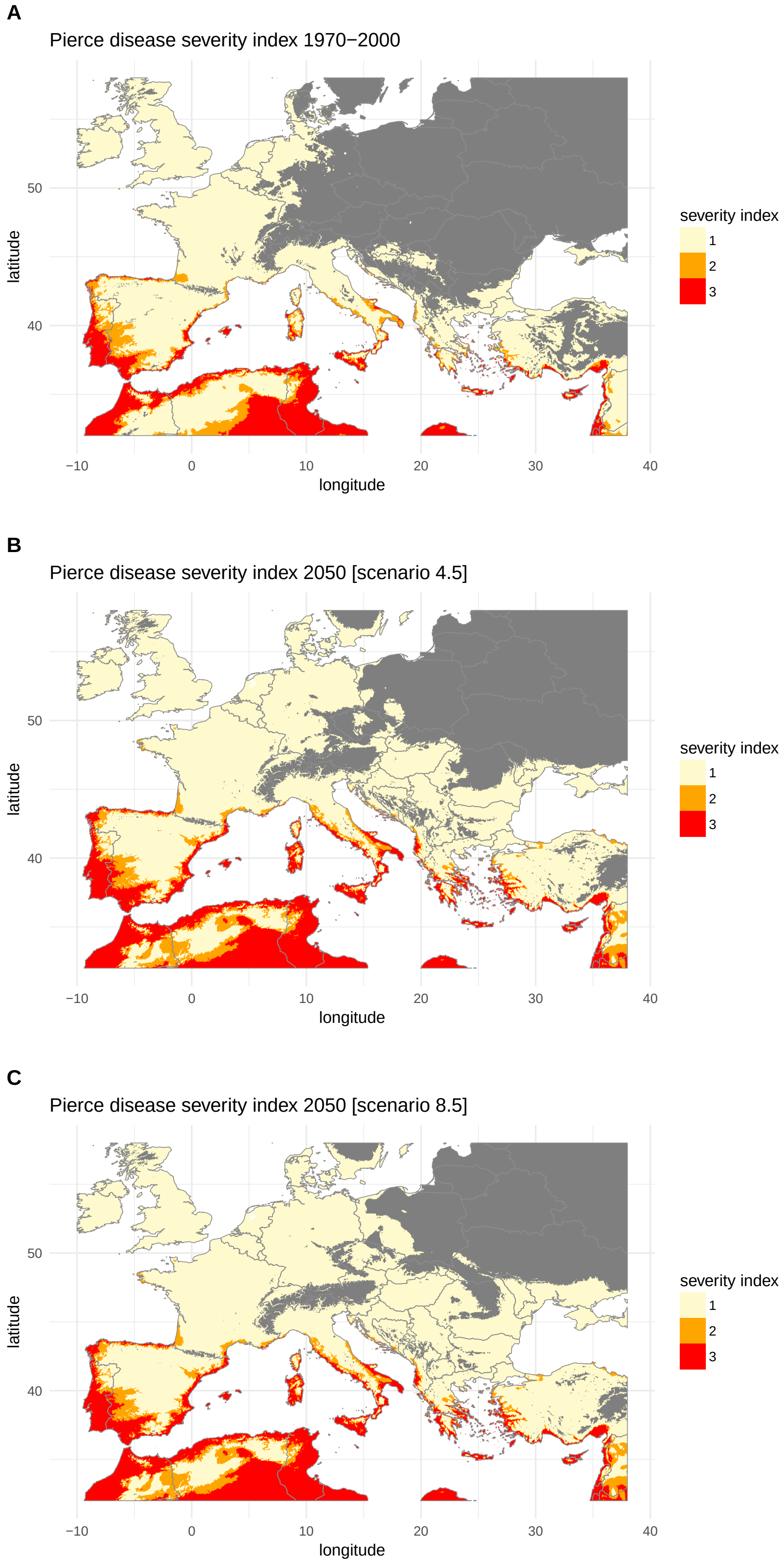

**Figure S5** – Predicted potential severity of Pierce's disease in Europe under current and future climate conditions obtained from a cumulative link model (CLM) fitted with bio10 and bio11 (Mean Temperature of Warmest Quarter and Mean Temperature of Coldest Quarter respectively). 1 = low severity, 2 = moderate severity, 3 = high severity. Current climate conditions are average temperature for the period 1970-2000 extracted from the Worldclim database (`http://worldclim.org/version2`). Future climate estimates were obtained from the MIROC5 global climate model (scenarios 4.5 and 8.5). A: Predicted PD severity index for the period 1970-2000. B: Predicted PD severity index in 2050 with the scenarios RCP4.5. C: Predicted PD severity index in 2050 with the scenarios RCP8.5. The predictions associated to localities with a negative MESS index were discarded and depicted in grey.

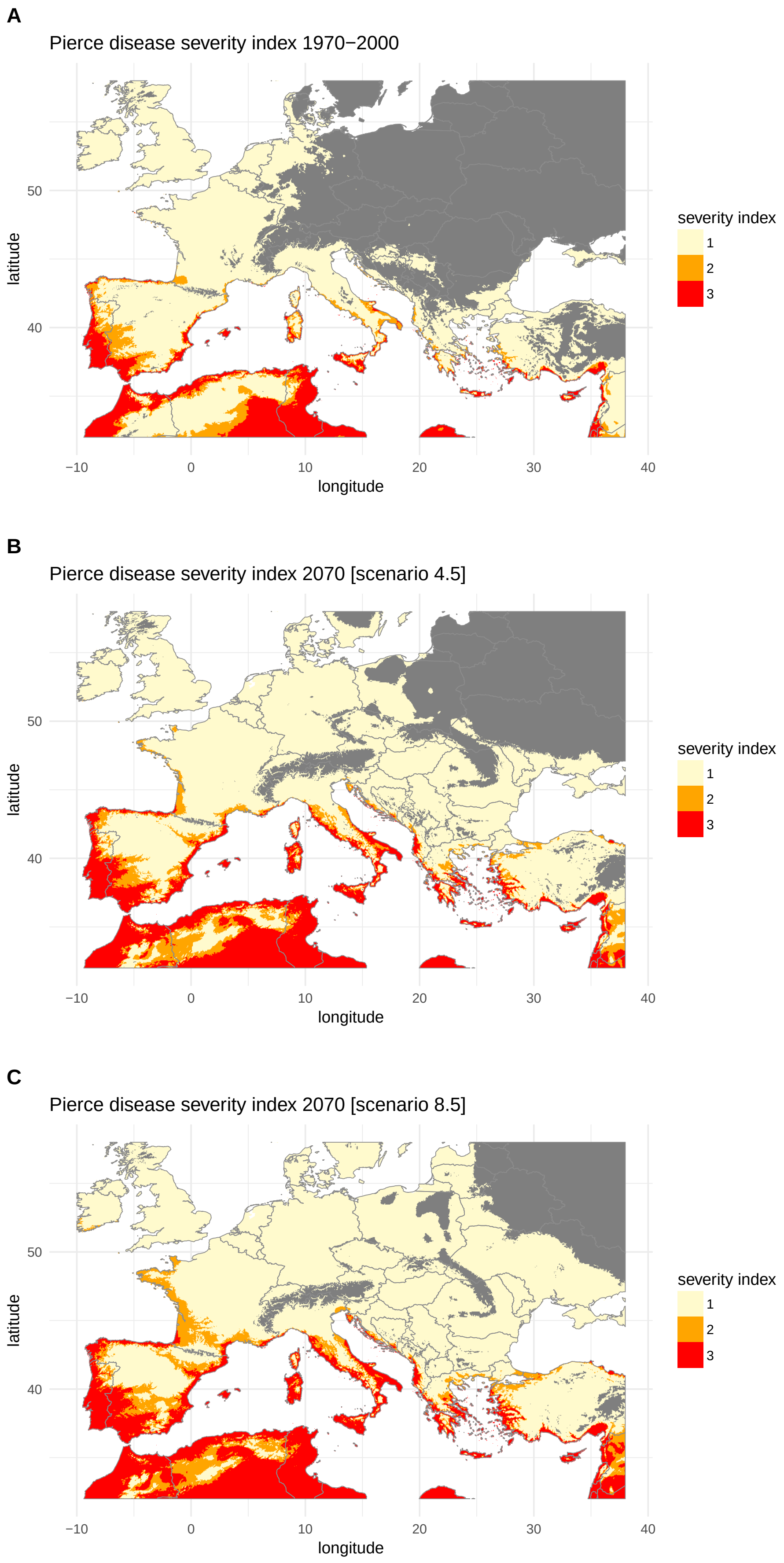

**Figure S6** – Predicted potential severity of Pierce's disease in Europe under current and future climate conditions obtained from a cumulative link model (CLM) fitted with bio10 and bio11 (Mean Temperature of Warmest Quarter and Mean Temperature of Coldest Quarter respectively). 1 = low severity, 2 = moderate severity, 3 = high severity. Current climate conditions are average temperature for the period 1970-2000 extracted from the Worldclim database (`http://worldclim.org/version2`). Future climate estimates were obtained from the MIROC5 global climate model (scenarios 4.5 and 8.5). A: Predicted PD severity index for the period 1970-2000. B: Predicted PD severity index in 2070 with the scenarios RCP4.5. C: Predicted PD severity index in 2070 with the scenarios RCP8.5. Areas associated to climate conditions that are not met within the range of conditions characterizing the set of reference points in the native range (i.e. MESS index $< 0$ see material and methods) are shown in grey.

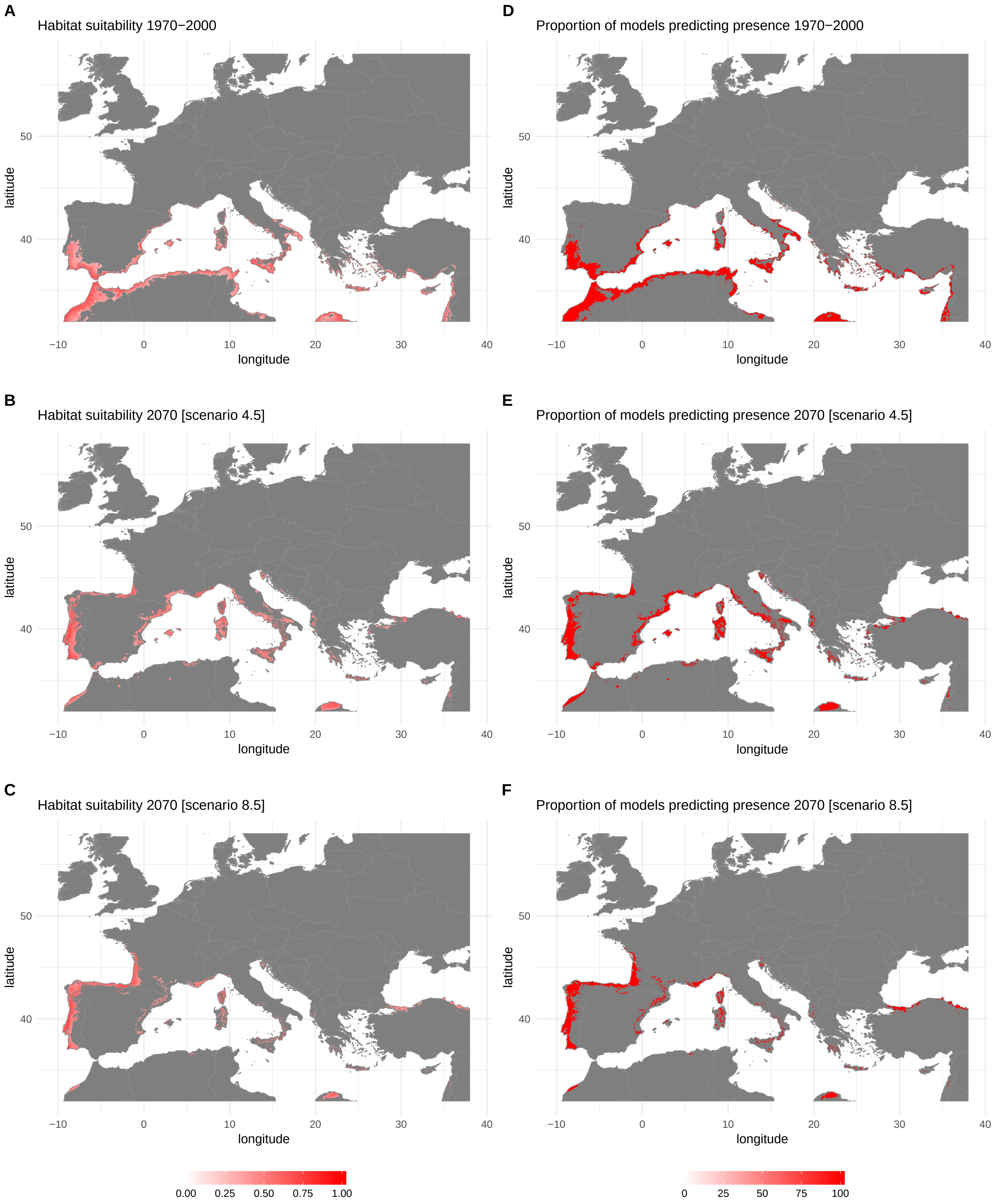

**Figure S7** – Predicted potential distribution of *Xylella fastidiosa* subsp. *pauca* in Europe under current and future climate conditions obtained by fitting Bioclim and Domain models. Current climate conditions are average temperatures for the period 1970-2000 extracted from the Worldclim database. Future climate estimates were obtained from the MIROC5 global climate model (scenarios 4.5 and 8.5). A: Habitat suitability for the period 1970-2000. B: Habitat suitability in 2070 for the scenario RCP4.5. C: Habitat suitability in 2070 for the scenario RCP8.5. D: Proportion of models predicting presence for the period 1970-2000. E: Proportion of models predicting presence in 2070 for the scenario RCP4.5. F: Proportion of models predicting presence in 2070 for the scenario RCP8.5. Maps A, B, C were obtained by averaging (ensemble forecasting) of the outputs of the models Bioclim and Domain run with 3 different climate datasets (see details in Table 1). Maps D, E, F were obtained by averaging the presence/absence maps derived from habitat suitability using the lowest presence threshold. Areas associated to climate conditions that are not met within the range of conditions characterizing the set of reference points in the native range (i.e. MESS index $< 0$ see material and methods) are shown in grey.

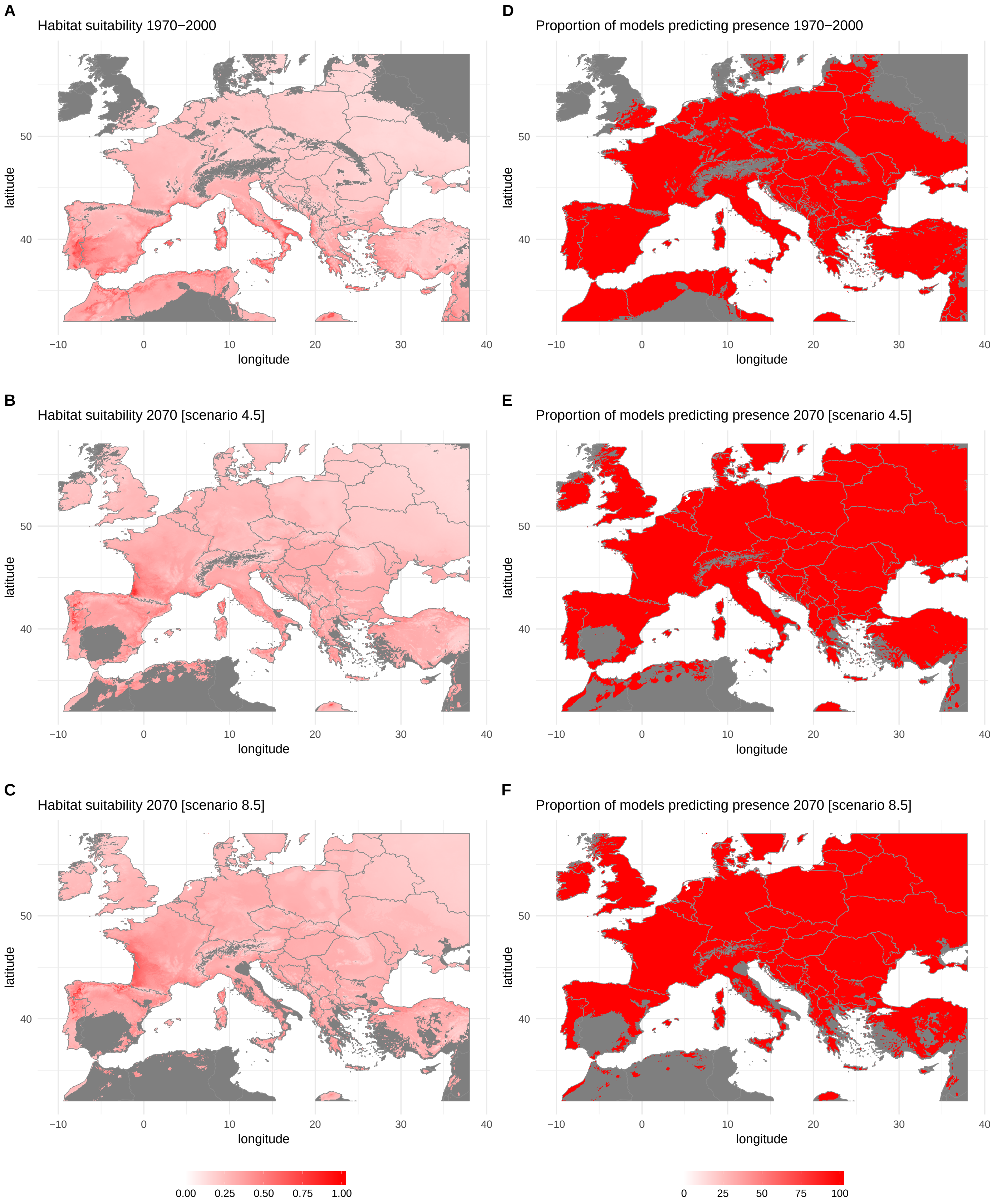

**Figure S8** – Predicted potential distribution of *Xylella fastidiosa* subsp. *multiplex* in Europe under current and future climate conditions obtained by fitting Bioclim and Domain models. Current climate conditions are average temperatures for the period 1970-2000 extracted from the Worldclim database. Future climate estimates were obtained from the MIROC5 global climate model (scenarios 4.5 and 8.5). A: Habitat suitability for the period 1970-2000. B: Habitat suitability in 2070 for the scenario RCP4.5. C: Habitat suitability in 2070 for the scenario RCP8.5. D: Proportion of models predicting presence for the period 1970-2000. E: Proportion of models predicting presence in 2070 for the scenario RCP4.5. F: Proportion of models predicting presence in 2070 for the scenario RCP8.5. Maps A, B, C were obtained by averaging (ensemble forecasting) of the outputs of the models Bioclim and Domain run with 3 different climate datasets (see details in Table 1). Maps D, E, F were obtained by averaging the presence/absence maps derived from habitat suitability using the lowest presence threshold. Areas associated to climate conditions that are not met within the range of conditions characterizing the set of reference points in the native range (i.e. MESS index < 0 see material and methods) are shown in grey.

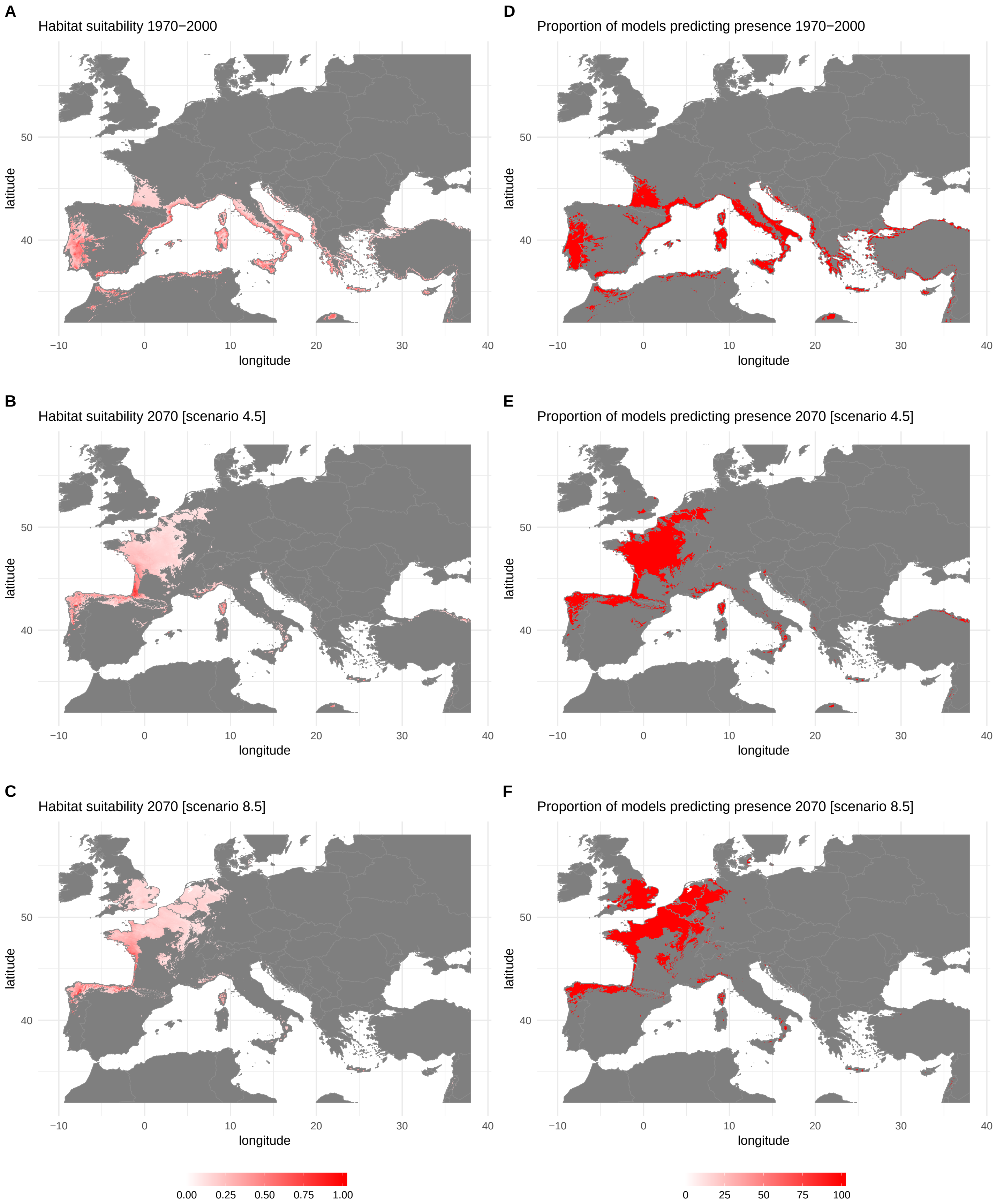

**Figure S9 –** Predicted potential distribution of *Xylella fastidiosa* subsp. *multiplex* ST6 & ST7 in Europe under current and future climate conditions obtained by fitting Bioclim and Domain models. Current climate conditions are average temperatures for the period 1970-2000 extracted from the Worldclim database. Future climate estimates were obtained from the MIROC5 global climate model (scenarios 4.5 and 8.5). A: Habitat suitability for the period 1970-2000. B: Habitat suitability in 2070 for the scenario RCP4.5. C: Habitat suitability in 2070 for the scenario RCP8.5. D: Proportion of models predicting presence for the period 1970-2000. E: Proportion of models predicting presence in 2070 for the scenario RCP4.5. F: Proportion of models predicting presence in 2070 for the scenario RCP8.5. Maps A, B, C were obtained by averaging (ensemble forecasting) of the outputs of the models Bioclim and Domain run with 3 different climate datasets (see details in Table 1). Maps D, E, F were obtained by averaging the presence/absence maps derived from habitat suitability using the lowest presence threshold. Areas associated to climate conditions that are not met within the range of conditions characterizing the set of reference points in the native range (i.e. MESS index $< 0$ see material and methods) are shown in grey.
